# Different functions of human scavenger receptors BI and BII overexpressed in a murine abdominal sepsis model

**DOI:** 10.64898/2026.02.04.703853

**Authors:** Naoki Hayase, Tatyana G. Vishnyakova, Irina N. Baranova, Alexander V. Bocharov, Xuzhen Hu, Amy P. Patterson, Peter S.T. Yuen, Thomas L. Eggerman, Robert A. Star

## Abstract

Class B scavenger receptors BI (SR-BI) and BII (SR-BII) internalize lipoproteins but also bind and internalize bacteria. Their roles in sepsis are unknown. We overexpressed human SR-BI and BII in the liver and kidney as well as bone marrow-derived macrophages, and then performed cecal ligation and puncture (CLP) surgery. SR-BI and BII transgenic mice had significantly worse survival compared to WT mice. 24 h after CLP, liver injury markers and histological damage were prominent in both SR-BI and BII transgenic mice, whereas kidney damage was similar. Systemic inflammatory cytokines were markedly increased in SR-BI and BII transgenic mice; parallel increases were seen in liver mRNA expression, not in the kidney. The highest degree of neutrophil infiltration was observed in the liver of SR-BI. Human SR-BI and BII dramatically decreased bacterial accumulation in the liver. Green fluorescence protein-labeled *E. coli* were efficiently phagocytosed in hepatic macrophages of SR-BI and BII transgenic mice; phagocytosis was more prominent in SR-BII transgenic mice. Finally, human SR-BI overexpression reduced systemic HDL-C level, eliminated adrenal cortex lipid droplets, and dampened the systemic increase of corticosterone after CLP. Supplementation with glucocorticoid and mineralocorticoid improved survival in SR-BI, but not SR-BII, transgenic mice after CLP. In summary, our findings suggest human SR-BI and BII overexpression contributes to higher mortality after CLP by excessive inflammatory response due to adrenal insufficiency (SR-BI) or hyperactive phagocytosis (SR-BII) in the liver.

## Introduction

Sepsis is a life-threatening condition characterized by excessive host immune response to infection. Sepsis-induced immune alterations have been associated with both poor short- and long-term outcomes^1^. These initial alterations are thought to be driven, in part, by the activation of pattern-recognition receptors (PRRs), which initiate inflammatory responses upon sensing pathogen-associated molecular patterns (PAMPs), leading to the immune system hyperactivation^2^. However, so far, the immunoadjuvant treatments by blocking PRRs (e.g., Eritoran and Resatorvid, both of which are Toll-like receptor-4 inhibitors) have failed to improve the 28-day mortality of septic patients^3, 4^. A comprehensive understanding of PRR functions not only in the innate immune system but also in other biological systems, including metabolic and endocrine systems, is essential for developing more targeted PRR-based therapies and minimizing off-target effects.

The class B scavenger receptor BI (SR-BI) and its splicing variant BII (SR-BII) are predominantly expressed in the liver, steroidogenic tissues, parenchymal epithelial cells in many organs, and phagocytic cells^5–9^. These receptors are well known for their roles in lipid metabolism, mediating both the selective uptake of high-density lipoprotein cholesterol (HDL-C) and the efflux of cholesterol from peripheral tissues via HDL^5, 10^. Our earlier *in vitro* studies demonstrated that both SR-BI and SR-BII function as PRRs by binding to and facilitating internalization of bacteria, and initiating inflammatory responses through interactions with bacterial components such as lipopolysaccharide (LPS), lipoteichoic acid, and the cytosolic protein GroEL^11, 12^.

To evaluate the roles of SR-BI and SR-BII in *in vivo* animal models of sepsis, SR-BI/BII knockout (KO) mice were initially studied, as the dual knock-out was easy to produce. SR-BI/BII KO mice had elevated systemic inflammatory cytokines and higher mortality after cecal ligation and puncture (CLP) surgery than wild-type (WT) controls. These findings suggest that SR-BI and BII may play a protective role against CLP-induced sepsis, although it is unclear if the two spice variants have similar or different roles. Septic SR-BI/BII KO mice had a profound defect in corticosterone production (adrenal insufficiency)^13^, which complicated the interpretation of the experimental results. The same study further reported that treatment with corticosterone alone (glucocorticoid only) did not improve survival in SR-BI/BII KO mice^13^. In contrast, when using mice treated with glucocorticoid and mineralocorticoid before CLP surgery^14^, we demonstrated that SR-BI/BII KO attenuated proinflammatory responses and improved survival. Thus, the survival outcomes of SR-BI/SR-BII KO mice after CLP depends on the details of adrenal replacement.

Our previous *in vitro* studies also suggested SR-B receptors promote pathogen internalization, escape from lysosomal killing, cytosolic proliferation, and amplified inflammation^12, 14–16^. Therefore, on the basis of both *in vitro* and *in vivo* results with steroid replacement, we suggested that SR-BI and BII may exert net detrimental effects in sepsis.

The SR-BI/BII KO mice developed adrenal insufficiency during sepsis because the lack of SR-BI/BII prevented HDL-C uptake into the adrenal gland, and decreased both gluco-and mineralocorticoid responses which then worsened sepsis. To better understand the role of SR-Bs in sepsis, we generated SR-BI and BII transgenic mice, with the transgene overexpressed mainly in the liver. The knock-in mice are easier to generate than specific SR-BI and BII knock-out mouse lines. We assumed that we could then evaluate the roles of SR-BI and BII in innate immunity during sepsis with less impact of confounding adrenal insufficiency^17^. This study aimed to elucidate the pathogenetic roles of human SR-BI and BII in the CLP-induced sepsis model fully treated with fluid, antibiotics, and analgesics – similar to the treatments critically ill patients would receive in an intensive care unit.

## Materials and Methods

### Animals

All animal experiments were conducted according to the National Institutes of Health Guide for the Care and Use of Laboratory Animals (U.S. Department of Health and Human Services Public Health Services, National Institutes of Health, Bethesda, Maryland; publication No. 85-23, 1985) and approved by the NIDDK Animal Care and Use Committee. We obtained 8-to 12-week-old male C57BL/J mice with an average weight of 20 to 25 g from the Jackson Laboratory (Bar Harbor, ME). We previously developed human SR-BI and human SR-BII transgenic mice using the liver-specific expression vector pLiv-11, which contains the human apoE promoter, on a C57BL/6J background.^17, 18^. The hepatic control region enabled transgenes to be expressed predominantly in hepatocytes^19^. A TaqMan PCR assay confirmed that human SR-BI and SR-BII expression was highest in the liver of SR-BI and SR-BII transgenic mice, respectively. The expression of human SR-B was also detectable in the kidney, lung, and spleen, but at 100 to1000 times lower extent than in the liver^17, 18^. Moreover, we reported human SR-BI and BII were highly expressed in bone marrow-derived macrophages in the respective transgenic mice.^17^ The mice were kept under specific pathogen-free conditions in a 12 h light/dark cycle with free access to chow and water, and acclimatized for at least 1 week before use. All surgeries were performed on a heated operating table under isoflurane anesthesia during the daytime. An extended-release formulation of buprenorphine (0.5 mg/kg Buprenorphine ER, SR Veterinary Technologies, Windsor, CO) was subcutaneously administered for analgesia immediately before the surgeries and every 72 hours for survival studies.

### Total RNA isolation and quantitative PCR analysis of proinflammatory cytokines in livers and kidneys

Collected liver and kidney samples were homogenized in TRIzol reagent using a Precellys 24 homogenizer (Bertin Technologies, Montigny-le-Bretonneux, France). RNA was isolated with the PureLink RNA mini kit (Thermo Fisher Scientific, Waltham, MA) after DNase treatment. Complementary DNA was synthesized from total RNA with a TaqMan reverse transcriptase reagent kit (Thermo Fisher Scientific). Quantitative reverse transcription polymerase chain reaction (RT-PCR) was performed with a StepOne real-time PCR system (Applied Biosystems, Foster City, CA). The mRNA levels were examined using TaqMan gene expression assays (Applied Biosystems, Foster City, CA) and TaqMan primers for chemokine (C-X-F motif) ligand 1 (CXCL1, Mm04207460_m1), tumor necrosis factor-α (TNF-α, Mm00443258_m1), interleukin-6 (IL-6, Mm00446190_m1), and glyceraldehyde-3-phosphate dehydrogenase (GAPDH, Mm03302249_m1). Relative levels of gene expression were calculated by the comparative cycle threshold (CT) method with GAPDH genes as a reference gene. All gene expression results were analyzed using the 2^-ΔΔCT^ method and are presented as normalized fold changes compared to the averaged results for corresponding sham-operated controls.

### Cecal ligation and puncture

Cecal ligation and puncture (CLP) was conducted as reported previously^20^. Briefly, a 4-0 silk ligature was placed 10 mm from the cecal tip. The cecum was punctured twice with a 21-gauge needle and gently squeezed to express a 1 mm column of fecal content. Prewarmed 0.9% saline (40 ml/kg) was injected intraperitoneally. Sham-operated mice were subjected to identical procedures except for the ligation and puncture of the cecum. Treatment with fluid and antibiotics was started at 6 h after surgery with subcutaneous injection of imipenem/cilastatin (14 mg/kg Primaxin, Merck, Whitehouse Station, NJ) in 40 ml/kg of 0.9% saline. In the acute study, animals were euthanized 24 h after surgery to collect blood and organ specimens. In the 7-day survival study, the treatment was continued every 12 h with imipenem/cilastatin (7 mg/kg) in 40 ml/kg of 0.6% saline. Additional doses of Buprenorphine ER were administered every 72 h after surgery. Sepsis severity was determined according to Murine Sepsis Score with minor modifications^21^. Animals exceeding a predefined threshold of the severity score (≥15) were euthanized and counted as non-survivors.

### E. coli injection model

*E. coli* K12 and green fluorescence protein (GFP)-labeled *E. coli* (ATCC, Manassas, VA) were grown, harvested, washed with phosphate-buffered saline (PBS) twice, and diluted to 1.6 × 10^7^ c.f.u./ml in PBS. To simulate the pathway of invading bacteria in the CLP sepsis model, mice were intraperitoneally injected with 600 μL of bacterial suspension and perfused with 20 ml PBS after 8 h. Then the liver and kidneys were harvested.

### Immunofluorescence analyses of mouse livers and kidneys

Harvested liver and kidneys were frozen in Tissue-Tek OCT compound (Sakura Finetek, Torrance, CA) and cryosectioned at a thickness of 5 μm. The sections were fixed with 4% paraformaldehyde (PFA) for 20 min and blocked with 1% bovine serum albumin and 1% goat serum albumin in phosphate-buffered saline for 1 h at room temperature. Subsequently, the tissues were incubated overnight at 4°C in 2 μg/ml rabbit anti-mouse F4/80 antibody (Bio-Rad, Hercules, CA) as a primary antibody. After washing, the sections were incubated with Alexa 647–goat anti-rabbit IgG (2 μg/ml) for 1 h at room temperature (Thermo Fisher Scientific). DNA was stained with 2 μg/ml Hoechst 33342 (Thermo Fisher Scientific). Confocal images were acquired using a Zeiss LSM 700 confocal microscope (Zeiss, Jena, Germany). The 3–6 randomly selected nonoverlapping fields were captured for the individual liver and kidney section using oil immersion objectives with ×20 or ×63 magnification. The total number of GFP-labeled bacteria was determined, and the proportion of bacteria localized inside F4/80-positive cells was then calculated. In a separate experiment, F4/80-positive cells were counted in each field and then normalized by the total nucleus count. The above-mentioned imaging analysis was conducted in a blinded manner using Fiji (National Institutes of Health, Bethesda, MD).

### Histological examination of organ damage and neutrophils

Liver and kidney specimens were fixed with 10% formalin and embedded in paraffin. The 4-μm sections were stained with periodic acid-Schiff (PAS) reagent. The extent of tubular injury was evaluated by the semiquantitative scoring system in 10 non-overlapping cortical fields under ×400 magnification and averaged per mouse. The percentage of glycogen-positive area in the whole surface area in the liver was measured using Fiji (National Institutes of Health) in 4–7 images taken for each mouse under ×200 magnification to assess liver function. For identifying neutrophils, the naphthol AS-D chloroacetate esterase kit (Millipore Sigma, Burlington, MA) was used for the liver. Neutrophils infiltrating the liver were counted in 7–12 non-overlapping fields under ×400 magnification. The evaluator was blinded to the origin of the samples until after histological quantitation.

### Immunohistochemical analysis for green fluorescence protein-labeled E. coli

Liver and kidney sections were deparaffinized and incubated in sodium citrate buffer (10 mM sodium citrate, 0.05% Tween 20, pH 6.0) for 20 min at 98 °C using a microwave. Endogenous peroxidase activity was blocked with 0.3% hydrogen peroxide in methyl alcohol for 15 min. After blocking with goat serum, the specimens were incubated overnight at 4 °C with 5 µg/mL rabbit anti-GFP antibody (ab290; Abcam, Cambridge, MA, USA). Horseradish peroxidase-conjugated goat anti-rabbit antibody (Agilent Dako, Santa Clara, CA, USA) was applied to sections and incubated for 1 h at room temperature. Sections were developed using 3,3′-diaminobenzidine tetrahydrochloride (Sigma-Aldrich, St. Louis, MO, USA), and then counterstained with hematoxylin.

### Oil Red O staining

Adrenal glands were embedded in Tissue-Tek OCT compounds (Sakura Finetek) and cryosectioned at a thickness of 5 μm. The sections were fixed in 4% paraformaldehyde for 10 min. After incubating with 60% isopropanol for 2 minutes, the slides were stained with a freshly prepared Oil Red O working solution for 15 minutes. Then, the slides were rinsed with 60% isopropanol followed by hematoxylin staining for 15 seconds. The slides were rinsed with distilled water before mounting a coverslip onto the slides with warmed glycerol gelatin. The percentage of lipid droplet area in the adrenal cortex was measured with Fiji (National Institutes of Health) under ×400 magnification.

### Measurement of blood chemistry, cytokines, vascular endothelial growth factor, high-density lipoprotein cholesterol, and corticosterone

Blood chemistry (serum blood urea nitrogen, aspartate aminotransferase, and alanine aminotransferase) was measured using commercial assays (VRL Diagnostics, Gaithersburg, MD). Interleukin-6 (IL-6), tumor necrosis factor-alpha (TNF-a), vascular endothelial growth factor (VEGF), high-density lipoprotein cholesterol (HDL-C) and corticosterone were measured using the corresponding mouse enzyme-linked immunosorbent assay (ELISA) kits according to the manufacturer’s instructions. IL-6, TNF-a, and VEGF ELISA kits were obtained from R&D Systems (Minneapolis, MN). HDL-C and corticosterone ELISA kits were purchased from AFG Scientific (Northbrook, IL) and Enzo Life Sciences (Farmington, NY), respectively.

### Bacterial count in blood, peritoneal lavage fluid, and organs

Anesthetized mice were washed with 70% ethanol under sterile condition at 24 h after CLP surgery. Peritoneal lavage was performed with 5 ml sterile PBS. The lavage fluid was collected using an 18-gauge needle before blood was drawn by cardiac puncture. The liver and kidneys were harvested and homogenized. Serial dilutions of blood, peritoneal lavage fluid, and organ homogenate were plated on 5% sheep blood agar plates (BD, Franklin Lakes, NJ). The plates were incubated at 37 °C for 24 h and bacterial colonies were counted. A value of 0.5 colony-forming unit was assigned to the samples that did not have detectable bacterial colonies.

### Glucocorticoid and mineralocorticoid supplementation

6α-methylprednisolone and fludrocortisone acetate were obtained from Sigma-Aldrich. A glucocorticoid and mineralocorticoid (GM) cocktail was prepared at a similar potency as described in the previous study.^22^ Mice were subcutaneously supplemented with vehicle (20 ml/kg of 3% dimethyl sulfoxide diluted with 0.9% saline) or a cocktail of 1.6 mg/kg 6α-methylprednisolone and 40 μg/kg fludrocortisone acetate in vehicle immediately after CLP surgery. The treatment with the vehicle or GM cocktail was continued daily at half doses for the following 4 days.

### Study design

We used three experimental groups for 7-day survival analysis, microbiological evaluation and immunofluorescence assays using GFP-labeled *E. coli*: Wild-type (WT), SR-BI, and SR-BII groups. Assessment of biochemistry, histological organ damage, inflammatory cytokines, and VEGF, HDL-C, corticosterone, and lipid droplets were conducted using six experimental groups: sham-operated or baseline WT, SR-BI, and SR-BII groups and CLP-injured WT, SR-BI, and SR-BII groups. Additionally, two groups were employed to investigate the impact of GM supplementation on septic SR-BI and BII transgenic mice: GM-treated and vehicle-treated groups. The experimental unit was a single animal.

### Randomization and blinding

The investigators were blinded to the genotype and treatment group in all experiments. A third-party allocated a random number to each mouse cage ID using the RAND function (Excel version 16, Microsoft, Redmond, WA). The IDs were then sorted from smallest to largest allocation number to create a randomly ordered ID list. Letter labels were assigned to the cage IDs in this order. The third party replaced the original cage card with the corresponding letter card. The surgeon (NH) performed CLP surgery, bacterial infusion, or a sham operation in alphabetical order of the letter cards. The WT, SR-BI, and BII transgenic mice had a similar appearance. The third-party randomized SR-BI and BII mice into the vehicle and GM treatments using the RAND function and prepared the drug for each mouse according to the allocated group.

### Sample size calculation

In a pilot study, we compared the 7-day survival rates of WT (n = 7), SR-BI transgenic (n = 8), and SR-BII transgenic mice (n = 9) after CLP surgery. The respective survival rates were 28.6, 0.0, and 0.0%. To detect a 29% difference in the survival rate in both comparisons: WT vs. SR-BI transgenic and WT vs. SR-BII transgenic, we estimated the effect size as 1.137 and calculated a sample size of 14 mice per group, assuming a two-sided alpha level of 0.05 and power of 0.80.

### Statistical analysis

The Shapiro–Wilk test was used to assess the normality of distribution of all continuous variables. The data are expressed as means ± SD when normally distributed and as medians (interquartile ranges) when not normally distributed. For normally distributed data, the Student’s t-test was applied to analyze a significant difference between groups. Otherwise, the Wilcoxon rank-sum test was used as a nonparametric test. The survival rates were represented using Kaplan-Meier survival curves and compared using a log-rank test. Multiple pairwise comparisons were conducted using a one-way ANOVA followed by the Holm–Šídák test for parametric values and a lognormal one-way ANOVA followed by Holm–Šídák test or the Kruskal–Wallis test followed by the Dunn’s test for nonparametric values. Repeated measures analysis was performed using a two-way ANOVA with the Tukey–Kramer test. A two-tailed P value of less than 0.05 was considered statistically significant for all tests. All calculations were conducted using GraphPad Prism 10 software (GraphPad Software Inc., La Jolla, CA).

## Results

### Overexpression of human SR-B transgenes worsened the survival and sepsis severity of septic mice

We conducted a 7-day survival study to examine whether overexpressed human SR-BI or BII could affect the survival of septic mice after CLP surgery. Both human SR-BI and SR-BII transgenic mice had a significantly lower survival rate than WT mice (SR-BI vs. WT, 6.3% vs. 33.3%, p = 0.001; SR-BII vs. WT, 6.3% vs. 33.3%, p = 0.001). The survival rates of those transgenic mice plummeted to 6.3% within 96 h after CLP, while that of WT mice decreased more gradually and was still 80% at 96 h. Additionally, SR-BI transgenic mice had a comparable survival rate to SR-BII transgenic mice (Figure 1A). Collectively, human SR-BI and SR-BII overexpression resulted in the lower 7-day survival of CLP-induced septic mice.

**Figure 1.**
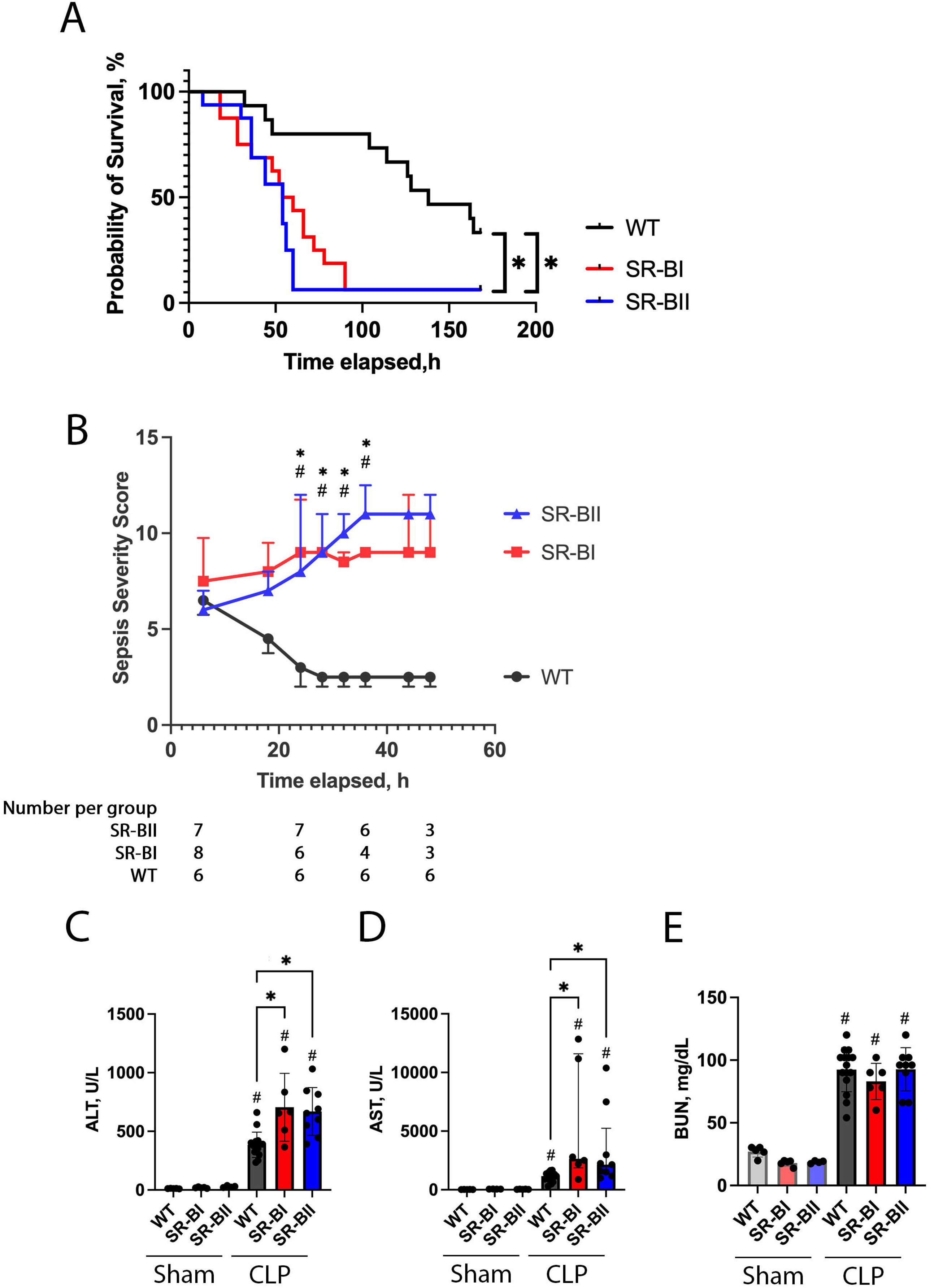
Kaplan-Meier survival curves, sepsis severity scores, circulating hepatic and renal injury markers after cecal ligation and puncture surgery. (A) A seven-day survival study was conducted for wild-type (WT) and human class B scavenger receptor BI (SR-BI) and BII (SR-BII) transgenic mice following cecal ligation and puncture (CLP) surgery. Kaplan-Meier survival curves are shown. SR-BI and BII transgenic mice had significantly worse survival compared to WT mice (SR-BI (red) vs. WT (black), 6.3% vs. 33.3%, p = 0.001; SR-BII (blue) vs. WT (black), 6.3% vs. 33.3%, p = 0.001; n = 15 in the WT and n = 16 in the SR-BI and SR-BII groups; n = 47 total). *P <0.05 versus respective control. (B) The longitudinal changes in sepsis severity scores were assessed until the number of remaining animals in either experimental group was reduced to three following CLP surgery. The number of remaining mice in each group at 6, 24, 36, and 48 h is shown under the graph. The data are represented as means ± SD. The comparison between the groups at each time point was performed Tukey multiple comparison test after two-way ANOVA test. #P <0.05 between SR-BI and WT, *P <0.05 between SR-BII and WT. (C) Blood was collected at 24 h after CLP or sham operation. Serum alanine aminotransferase (ALT), aspartate aminotransferase (AST), and blood urea nitrogen (BUN) levels were measured and compared between the six experimental groups (sham WT, n = 5; sham SR-BI, n = 5; sham SR-BII, n = 4; CLP WT, n = 5; CLP SR-BI, n = 5; CLP SR-BII, n = 5; n = 44 total). ALT and BUN data are shown as means ± SD due to their normal distribution. The data of AST are represented as medians (interquartile ranges) due to their skewed distribution. #P <0.05 versus sham, *P <0.05 between WT, SR-BI and SR-BII after CLP or sham.

The sepsis severity score of SR-BI and BII transgenic mice was significantly higher than that of WT mice at 24 h after CLP (Figure 1B). The survival curves began to diverge at 24 h. More than 20% of SR-BI transgenic septic mice died or were euthanized after 24 h (Figure 1A). Therefore, we chose 24 h after CLP as the best timepoint to study the underlying mechanisms.

### Overexpression of human SR-B transgenes exacerbated sepsis-induced liver injury

Next, we asked why overexpressed human SR-BI and BII increased the mortality of septic mice. Our previous study confirmed higher mRNA expression and protein production of human SR-B in liver and kidney of SR-B transgenic mice compared to WT mice^17^. Intraperitoneal injection of lipopolysaccharide caused more severe histological damage and immune response in the liver and kidney of SR-B transgenic mice^17^. Therefore, we focused on the liver and kidney, hypothesizing that organ damage could be involved in higher mortality during CLP sepsis.

CLP sepsis caused the elevation of liver and kidney injury markers (ALT, AST, and BUN) in WT and SR-B transgenic mice at 24 h. ALT and AST were significantly increased in SR-BI (75% and 127% increase in ALT and AST, respectively) and SR-BII transgenic mice (73% and 84% increase in ALT and AST, respectively) vs. WT mice. In contrast, the BUN level was similar across the experimental groups (Figure 1C).

We then evaluated the histological liver and kidney damage at 24 h after CLP. As expected, CLP sepsis decreased liver glycogen storage represented by deep pink region in PAS staining. Liver glycogen was additionally and significantly decreased in SR-BI and BII transgenic mice (84% and 90% reduction in SR-BI and BII, respectively, vs. WT septic mice, Figure 2A and C). All the experimental groups developed similar sepsis-induced kidney tubular damage as characterized by increased vacuolization and loss of brush border to a similar extent (Figure 2B and D).

**Figure 2.**
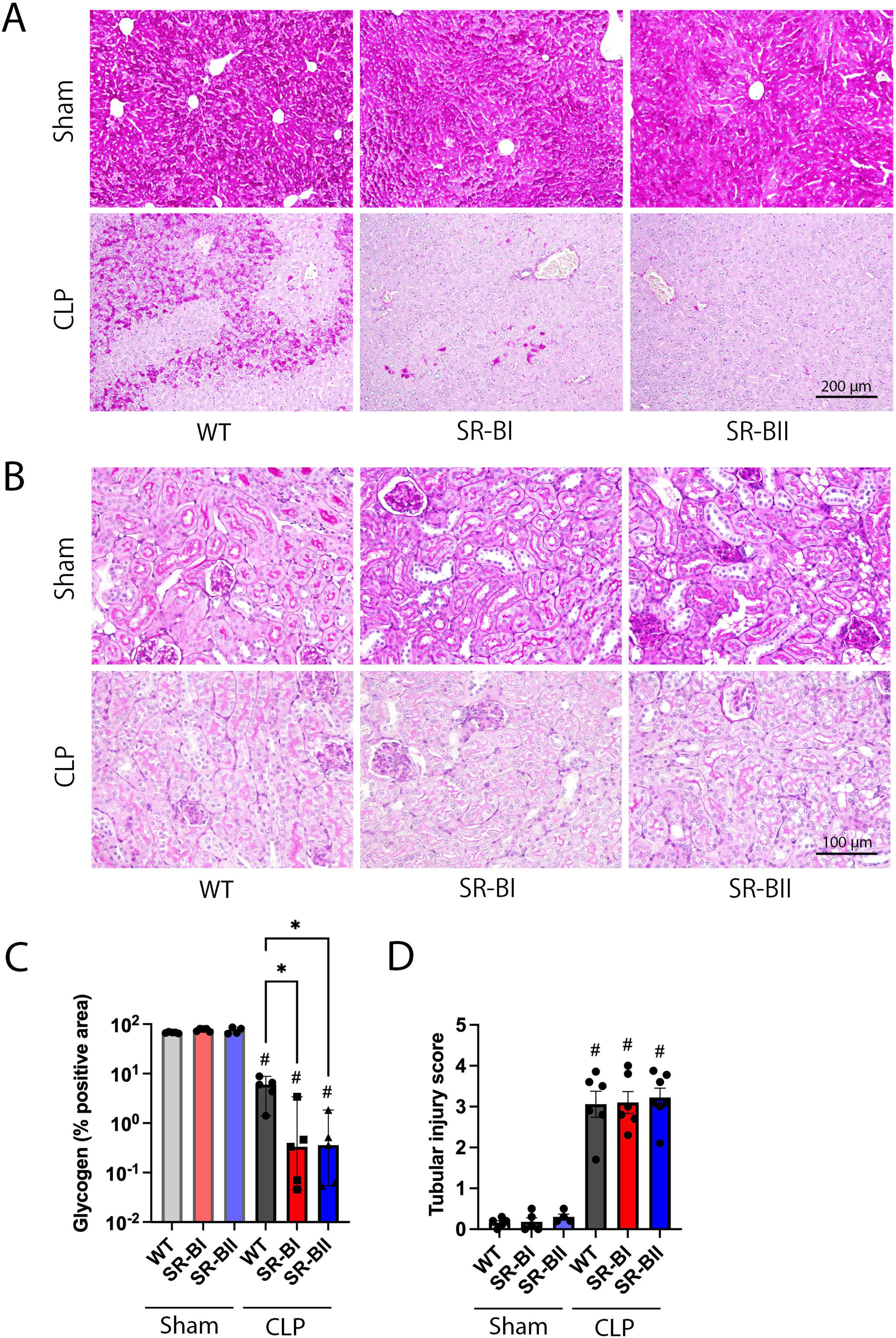
Liver and kidney histological damage at 24 h after cecal ligation and puncture surgery. The liver and kidneys were harvested from Wild-type (WT) and human class B scavenger receptor BI (SR-BI) and BII (SR-BII) transgenic mice at 24 h after cecal ligation and puncture (CLP) surgery. (A) Representative images of Periodic acid-Shiff (PAS) staining in the liver are shown. The dark pink area represents liver glycogen, which was significantly reduced after CLP. (B) Representative images of PAS staining in the renal cortex are shown. CLP induced distinct kidney damage characterized by vacuolar degeneration and blush border loss. (C) The percentage of glycogen-rich area in whole tissue area was measured using Fiji and compared among the experimental groups (sham WT, n = 5; sham SR-BI, n = 5; sham SR-BII, n = 4; CLP WT, SR-BI, and SR-BII, n = 5 per group; n = 29 total). The data are represented as medians (interquartile ranges). #P <0.05 versus sham, *P <0.05 between WT, SR-BI and SR-BII after CLP or sham. (D)The extent of kidney damage was assessed using tubular injury scores and compared across the experimental groups (sham WT, n = 5; sham SR-BI, n = 5; sham SR-BII, n = 4; CLP WT, SR-BI, and SR-BII n = 7 per group; CLP SR-BII, n = 7; n = 44 total). The data are shown as means ± SD. #P <0.05 versus sham, *P <0.05 between WT, SR-BI and SR-BII after CLP or sham.

Taken together, both human SR-BI and BII transgenic mice developed significantly more liver injury compared to WT mice during CLP sepsis despite similar levels of kidney injury.

### Human SR-B enhanced systemic and hepatic proinflammatory response; neutrophil infiltration was prominent in the liver of SR-BI transgenic mice

Our previous *in vivo* study using the LPS injection model reported that the production of proinflammatory cytokines and chemokines mediated through the mitogen-activated protein kinase (MAPK) signaling pathway might be a driver of organ injury in human SR-B transgenic mice with endotoxemia^17^. Thus, we evaluated systemic and organ levels of proinflammatory cytokines using ELISA and quantitative RT-PCR. As expected, plasma IL-6 and TNF-α levels were dramatically increased after CLP. Of note, those levels were more prominent in septic SR-B transgenic mice (IL-6, 28- and 12-fold higher in SR-BI and BII, respectively; TNF-α, 3.4- and 2.4-fold higher in SR-BI and BII, respectively; Figure 3A). Additionally, the PCR assay of inflammatory genes revealed that the expression levels of CXCL1, IL-6, and TNF-α in the liver were significantly increased in both SR-BI and BII transgenic mice at 24 h after CLP. The increase in the expression levels was larger than in WT (CXCL1, 5.7- and 8.0-fold higher in SR-BI and BII, respectively; IL-6, 5.3- and 4.0-fold higher in SR-BI and BII, respectively; TNF-α, 3.3- and 2.6-fold higher in SR-BI and BII, respectively; Figure 3B). In contrast, renal CXCL1, IL-6, and TNF-α mRNA levels were comparably elevated among the experimental groups after CLP surgery, although the CXCL1 level was more increased in SR-BII transgenic mice compared to WT (Figure 3C). Next, we hypothesized hyperinflammatory status of SR-B transgenic mice during sepsis could impair vascular endothelial integrity and increase vascular permeability. A marker of systemic vascular endothelial dysfunction, VEGF, was assessed. The plasma VEGF level was elevated at 24 h after CLP. The level was significantly higher in SR-B transgenic mice than in WT (61% and 49% higher in SR-BI and BII, respectively; Figure 3D).

**Figure 3.**
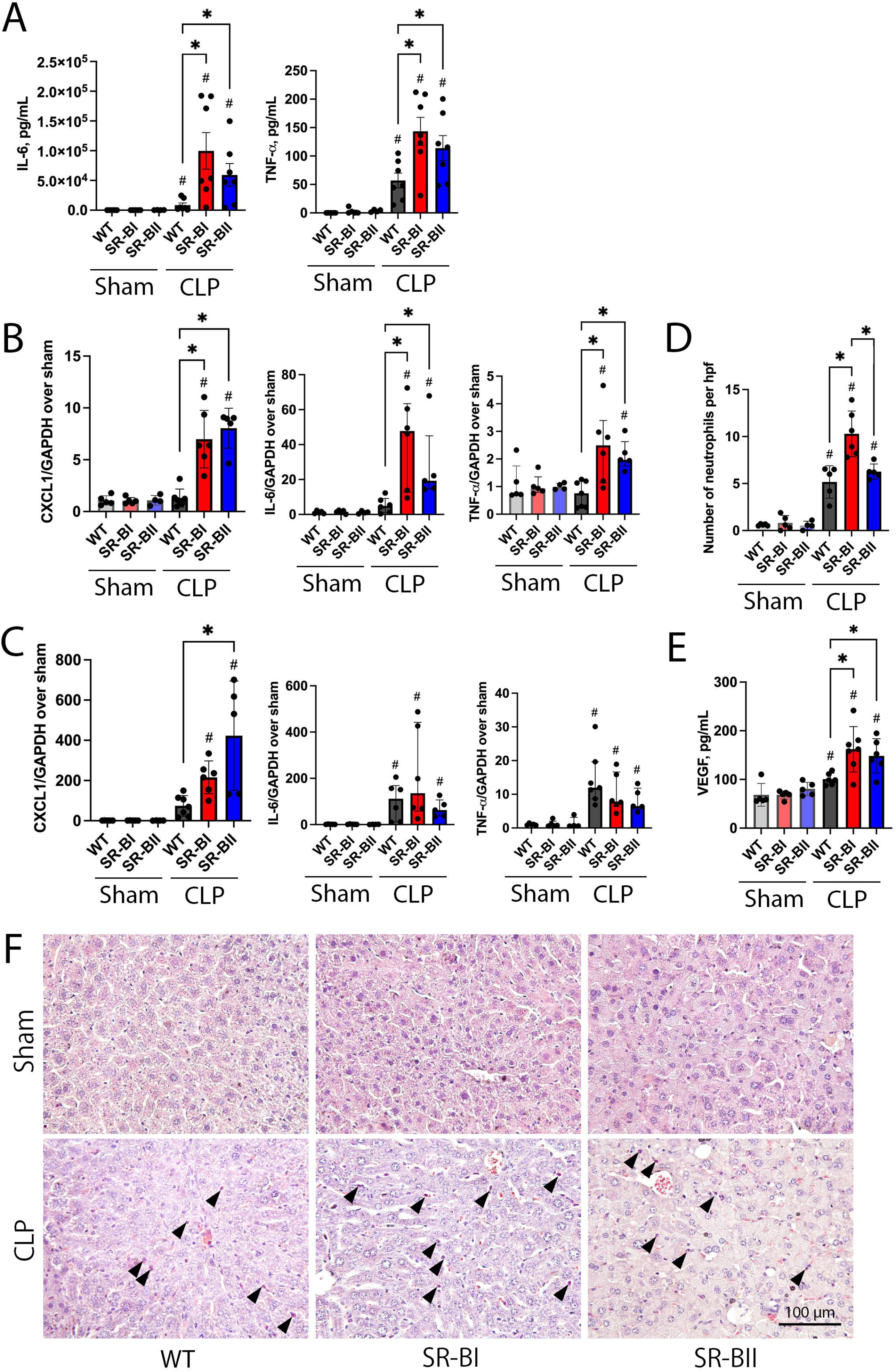
Inflammatory cytokines, vascular endothelial growth factor, and neutrophil infiltration at 24 h after cecal ligation and puncture. The blood, liver, and kidneys were collected from Wild-type (WT) and human class B scavenger receptor BI (SR-BI) and BII (SR-BII) transgenic mice at 24 h after cecal ligation and puncture (CLP) surgery. (A) Plasma levels of interleukin-6 (IL-6), tumor necrosis factor-α (TNF-α) were assessed and compared among the experimental groups (sham WT, n = 5; sham SR-BI, n = 5; sham SR-BII, n = 4; CLP WT, SR-BI, and SR-BII, n = 7 per group; n = 35 total). The data are shown as means ± SD. #P <0.05 versus sham, *P <0.05 between WT, SR-BI and SR-BII after CLP or sham. The mRNA expression levels of chemokine (C-X-F motif) ligand 1 (CXCL1), interleukin-6 (IL-6), and tumor necrosis factor-α (TNF-α) in the liver (B) and kidney (C) were evaluated and normalized with glyceraldehyde-3-phosphate dehydrogenase (GAPDH) in corresponding experimental groups for respective organs (sham WT, n = 5; sham SR-BI, n = 5; sham SR-BII, n = 4; CLP WT, n = 7; CLP SR-BI, n = 6 and SR-BII n = 5; n = 32 total). The CXCL1 expression levels are expressed as means ± SD due to their normal distribution, whereas IL-6 and TNF-α levels are expressed as medians (interquartile ranges) due to their skewed distribution. #P <0.05 versus sham, *P <0.05 between WT, SR-BI and SR-BII after CLP or sham. (D) Vascular endothelial growth factor (VEGF) was measured and compared among the experimental groups (sham WT, SR-BI, and SR-BII, n = 5 per group; CLP WT, n = 6; CLP SR-BI, n = 7 and SR-BII n = 6; n = 34 total). The data are shown as means ± SD. #P <0.05 versus sham, *P <0.05 between WT, SR-BI and SR-BII after CLP or sham. (E) Representative images of naphthol AS-D chloroacetate esterase staining in the liver are exhibited. The arrowheads denote infiltrating neutrophils. (F) The number of neutrophils per high-power field was compared among the experimental groups (sham WT and SR-BI, n = 5 per group; sham SR-BII, n = 4; CLP WT, n = 5; CLP SR-BI, n = 6; CLP SR-BII, n = 5; n = 30 total). The data are shown as means ± SD. #P <0.05 versus sham, *P <0.05 between WT, SR-BI and SR-BII after CLP or sham.

The naphthol AS-D chloroacetate esterase staining showed abundant liver neutrophil infiltration at 24 h after CLP. Liver from SR-BI transgenic mice had the highest infiltration of neutrophils among the experimental groups (Figures 3E and F).

Based on the above results, the overexpression of human SR-B transgenes exacerbated the inflammatory response, especially in the liver, and increased a marker of vascular endothelial dysfunction during CLP sepsis. The neutrophil infiltration was most pronounced in the liver of SR-BI transgenic mice.

### Human SR-B significantly reduced bacteria accumulation in the liver via promoting macrophage phagocytosis of bacteria; the phagocytosis was prominent in SR-BII transgenic mice

In our previous *in vitro* study using Hela cells overexpressing human SR-Bs, we suggested that both SR-BI and BII acted as pattern recognition receptors that promoted bacterial adhesion, internalization, cytosolic invasion, and intracellular survival and proliferation^12^. Therefore, we speculated SR-BI and BII transgenic mice would have increased bacterial invasion in distant organs in CLP sepsis.

Unexpectedly, we observed a decreasing trend in bacterial counts in the blood and peritoneal lavage fluid of septic SR-B transgenic mice at 24 h after CLP. This tendency was more pronounced in SR-BII transgenic mice, which had a 96% lower blood bacterial count compared to WT mice (Figure 4A and B). Of note, bacteria were nearly undetectable in the liver and kidney of SR-BI and BII transgenic mice during CLP-induced sepsis, despite abundant bacterial colonies detected in those organs of WT mice (Figure 4C and D).

**Figure 4:**
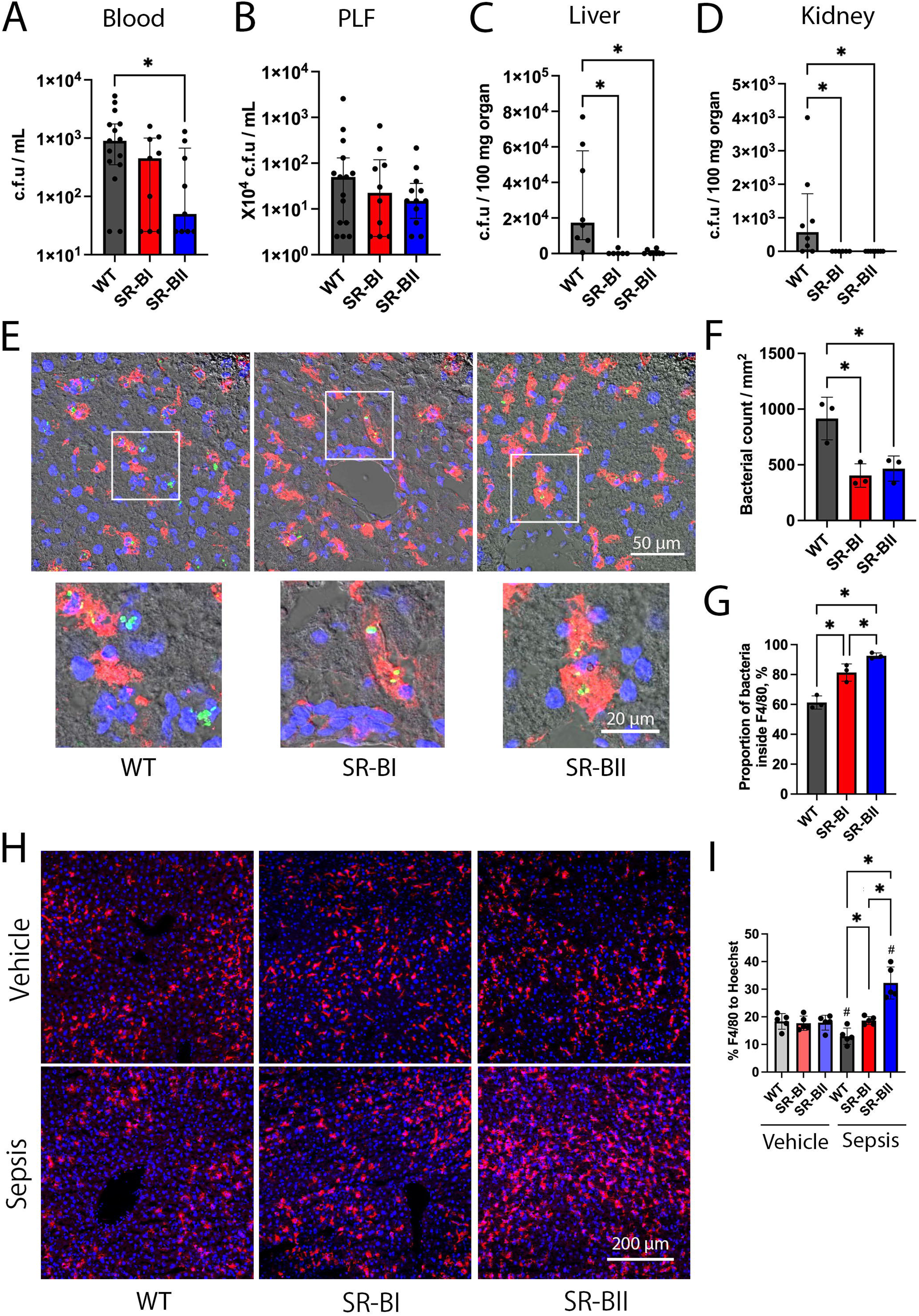
Bacterial counts after cecal ligation and puncture, and green fluorescence protein-labeled E. coli engulfed by macrophages after bacterial injection. The bacterial counts in the following specimens were examined and compared between Wild-type (WT) and human class B scavenger receptor BI (SR-BI) and BII (SR-BII) transgenic mice at 24 h following cecal ligation and puncture (CLP); (A) Blood (WT, n = 15; SR-BI, n = 9; SR-BII, n = 9; n = 33 total). (B) Peritoneal lavage fluid (PLF) (WT, n = 15; SR-BI, n = 10; SR-BII, n = 12; n = 37 total). (C) Liver and (D) kidney (WT, n = 8; SR-BI, n = 6; SR-BII, n = 8; n = 22 total). The data are shown as medians (interquartile ranges). #P <0.05 versus sham, *P <0.05 between WT, SR-BI and SR-BII. The immunofluorescence analyses were performed to observe localization of green fluorescence protein (GFP)-labeled *E. coli* and macrophages in liver harvested at 8 h after intraperitoneal bacterial infusion. (E) The representative images for liver specimens are shown. Positive staining for *E. coli*, F4/80, and DNA are indicated in green, red, and blue, respectively. The GFP-labeled bacteria were frequently localized in F4/80-positive cells in liver of SR-BI and BII transgenic mice. (F) The total number of bacteria per high-power field and (G) the percentage of bacteria localized within macrophages were averaged and compared across the experimental groups (WT, n = 3; SR-BI, n = 3; SR-BII, n = 3; n = 9 total). The data are represented as means ± SD. #P <0.05 versus sham, *P <0.05 between WT, SR-BI and SR-BII. Hepatic macrophages of Wild-type (WT) and human class B scavenger receptor BI (SR-BI) and BII (SR-BII) transgenic mice were observed under a confocal microscope at 8 h after intraperitoneal injection with vehicle or *E. coli* K12. (H) Representative images are shown. Positive staining for F4/80 and DNA is indicated in red and blue, respectively. (I) The percentage of F4/80-positive cells relative to the total number of nuclei was averaged for each mouse and compared across experimental groups (n = 5 per group; n = 30 total). The data are shown as means ± SD. #P <0.05 versus vehicle, *P <0.05 between WT, SR-BI and SR-BII after bacterial or vehicle infusion.

Next, we investigated the mechanism by which invading bacteria were eliminated in the liver and kidney. GFP-labeled *E. coli* were intraperitoneally injected to mimic enteric bacteria released into the peritoneal cavity. We previously detected higher levels of mRNA expression and protein production of human SR-BI and BII in bone marrow-derived macrophages (BMDMs) from SR-BI and BII transgenic mice compared to WT^17^. Resident and monocyte-derived macrophages were visualized with an immunofluorescence assay using anti-F4/80 antibody. We found an increased number of GFP-expressing bacteria exclusively localized within F4/80-positive cells of liver tissue harvested at 8 h after injection (Figure 4E). The total number of bacteria was 56% and 49% lower in the liver of SR-BI and BII transgenic mice, respectively, compared to that in the liver of WT mice, consistent with the liver bacterial counts after CLP surgery (Figure 4F).

Furthermore, the proportion of bacteria located within F4/80-positive cells relative to the total number of identified bacteria was 33% and 51% higher in SR-BI and BII transgenic mice, respectively, compared to WT. Bacteria were more frequently detected within the macrophages in SR-BII compared to SR-BI transgenic mice (Figure 4G). The kidney sections of all the experimental groups had too few GFP-labeled bacteria to perform analyses (Supplemental Figure 1).

Moreover, 4/80-positive cells were significantly decreased in the liver of WT mice at 8 h after *E. coli* injection (29% decrease from the baseline), whereas their number was unchanged in SR-BI transgenic mice, and markedly increased (80% increase from the baseline) in SR-BII transgenic mice (Figure 4H and I). This result indicates that extensive recruitment of monocyte-derived macrophages occurs in the liver of SR-BII transgenic mice.

According to several reports, SR-BI is abundantly expressed in hepatocytes and kidney epithelial cells, suggesting that those cells might internalize bacteria via the scavenger receptors^23, 24^.

Immunohistochemical analysis for GFP-labeled bacteria revealed that those bacteria were frequently located in sinusoids between hepatocytes, not within hepatocytes, at 8 h after injection (Supplemental Figure 2A). A few bacteria were observed in peritubular capillaries of kidneys isolated from all the experimental groups (Supplemental Figure 2B).

In summary, the results suggest that overexpression of human SR-BI and BII dramatically decreased bacteria accumulation in the liver due to increased phagocytosis and the number of macrophages compared to WT. This effect was larger in SR-BII transgenic mice.

### Overexpression of human SR-BI reduced systemic HDL-C level, storage of lipid droplets in the adrenal gland, and dampened the increase of circulating corticosterone in response to CLP sepsis

Both SR-BI and BII mediate HDL-C uptake^5, 10^. We previously reported that human SR-BI and BII transgenic mice exhibited significantly lower levels of total cholesterol compared to WT under physiological conditions. Both of the transgenic mice similarly increased adrenal glucocorticoid production after LPS injection; however, this increase was less than in WT mice ^17^. We assessed the effect of enhanced human SR-BI and BII expression on HDL-C and corticosterone levels following CLP. At baseline and 24 hours after CLP, plasma HDL-C levels were significantly lower in SR-BI transgenic mice than in WT mice (22% lower at both baseline and after CLP), probably because of increased HDL-C uptake in organs overexpressing SR-BI such as the liver. HDL-C levels in SR-BII transgenic mice were not as low as those in SR-BI transgenic mice (Figure 5A). Circulating corticosterone was increased in all experimental groups after CLP. However, the corticosterone levels in septic SR-BI transgenic mice were significantly lower than those in WT (39% reduction). The difference in the corticosterone levels between SR-BII and WT mice did not reach statistical significance (Figure 5B).

**Figure 5:**
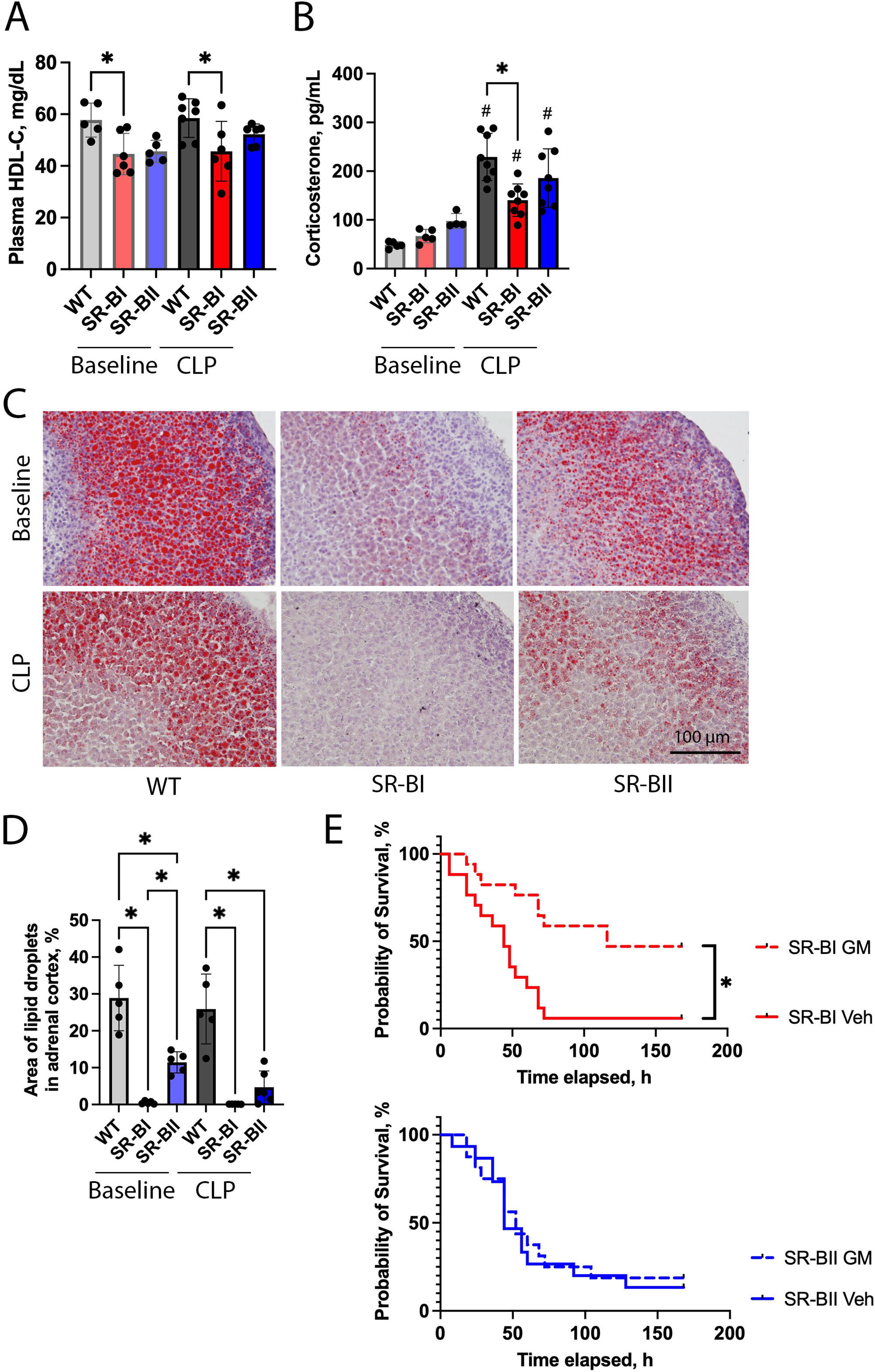
Systemic high-density lipoprotein cholesterol and corticosterone levels and lipid droplets in adrenal cortex at baseline and in sepsis. (A) High-density lipoprotein cholesterol (HDL-C) in the plasma of Wild-type (WT) and human class B scavenger receptor BI (SR-BI) and BII (SR-BII) transgenic mice were assessed at baseline or at 24 h after cecal ligation and puncture (CLP) surgery and compared among the experimental groups (baseline WT, n = 5; baseline SR-BI, n = 6; baseline SR-BII, n = 5; CLP WT, n = 7; CLP SR-BI, n = 6; CLP SR-BII, n = 6; n = 35 total). The data are shown as means ± SD. *P <0.05 between WT, SR-BI and SR-BII after CLP or at baseline. (B) The serum corticosterone levels were also evaluated at baseline or in CLP-induced sepsis (baseline WT, n = 5; baseline SR-BI, n = 5; baseline SR-BII, n = 4; CLP WT, SR-BI, and SR-BII, n = 8 per group; n = 38 total). The data are shown as means ± SD. #P <0.05 versus baseline, *P <0.05 between WT, SR-BI and SR-BII after CLP or at baseline. The lipid droplets in the adrenal cortex were examined using Oil-red O staining for the adrenal glands of WT and SR-BI and SR-BII transgenic mice at baseline or at 24 h after CLP. (C) The representative images for all the experimental groups were shown. The red area represents lipid droplets. (D) The percentage of lipid droplet area in the adrenal cortex was calculated using Fiji and compared across the groups (baseline WT, SR-BI, SR-BII, n = 5 per group; CLP WT, SR-BI, and SR-BII, n = 5 per group; n = 30 total). The data are shown as means ± SD. #P <0.05 versus sham, *P <0.05 between WT, SR-BI and SR-BII after CLP or at baseline. (D) A seven-day survival study was conducted for vehicle (Veh) and glucocorticoid and mineralocorticoid (GM) treatments in human class B scavenger receptor BI (SR-BI) and BII (SR-BII) transgenic mice following cecal ligation and puncture (CLP) surgery. Kaplan-Meier survival curves are shown. GM treatment significantly improved the survival rate of septic SR-BI (GM treatment (dotted red line) vs. vehicle treatment (solid red line), 47.1% vs. 5.9%, p < 0.001; n = 17 per group; n = 34 total), whereas it did not improve the survival in SR-BII transgenic mice (GM treatment (dotted blue line) vs. vehicle treatment (solid blue line), 18.6% vs. 13.3%, p = 0.81; n = 16 in the GM and n = 15 in the vehicle groups; n = 31 total). *P <0.05 versus respective control.

Cholesterol ester stored in adrenal gland lipid droplets provides an available pool of free cholesterol for corticosterone synthesis (functional reserve)^25^. We measured the area of lipid droplets in adrenal glands at baseline and following CLP surgery to examine this functional reserve. At baseline, both SR-BI and BII transgenic mice had significantly fewer lipid droplets in the adrenal cortex than WT (99% and 60% lower in SR-BI and BII, respectively), with lipid storage being particularly low in SR-BI transgenic mice. After CLP, lipid area remained stable in WT, decreased in SR-BII mice, and declined to nearly zero in SR-BI mice (Figure 5C and D).

To test whether the partial adrenal insufficiency contributed to higher mortality in septic SR-BI mice compared with WT, we conducted a 7-day survival study for vehicle and glucocorticoid and mineralocorticoid (GM) supplementation in SR-BI and BII transgenic mice after CLP. The GM treatment improved survival in SR-BI transgenic mice (GM treatment vs. vehicle treatment, 47.1% vs. 5.9%, p < 0.001), but not in SR-BII transgenic mice (GM treatment vs. vehicle treatment, 18.6% vs. 13.3%, p = 0.81) (Figure 5E).

Taken together, overexpressed human SR-BI remarkably reduced systemic HDL-C level, storage of lipid droplets in the adrenal gland, and dampened systemic increase of corticosterone in response to septic insult, while human SR-BII transgenic mice showed milder changes in these parameters. The increased mortality of SR-BI transgenic mice during CLP sepsis was mainly attributable to the partial adrenal insufficiency.

## Discussion

The dysregulated host immune response during sepsis causes poor clinical outcomes in septic patients^1^. Class B scavenger receptors BI and BII are involved in innate immunity, lipid metabolism, and hormonal response^5, 10, 11^. When studied *in vitro,* human SR-BI and BII act as pattern-recognition receptors to mediate uptake of gram-negative bacteria into cells^11, 12, 14^. To examine the role of those receptors in animal sepsis models, global SR-BI/SR-BII knock-out (KO) mice have been employed^13, 26^; although these studies were difficult to interpret because of multiple interacting abnormalities [excessive lymphocyte apoptosis, adrenal insufficiency (inhibition of HDL-C uptake, impaired accumulation of cholesterol esters), and female infertility^27, 28^.] For example, survival depended strongly on specific regimens of adrenal replacement. Furthermore, the SR-BI/SR-BII KO mice cannot detect SR-BI and BII-specific functions. Therefore, we developed human SR-B transgenic mice, with the transgenes expressed in liver and kidney as well as bone marrow-derived macrophages (BMDMs), and hoped this would help unravel their role in sepsis. We found: 1) Human SR-BI and SR-BII overexpression led to similar poor survival, liver damage, enhanced inflammatory responses, and decreased bacterial accumulation after CLP surgery compared to WT mice. 2) However, the mechanisms were different. SR-BII transgenic mice had higher bacterial phagocytic activity and a greater abundance of hepatic macrophages. In contrast, SR-BI transgenic mice had enhanced hepatic neutrophil infiltration. 3) Human SR-BI transgenic mice had partial adrenal insufficiency following sepsis; supplementation with glucocorticoid and mineralocorticoid (GM) reversed the survival deficit of septic SR-BI (but not SR-BII) transgenic mice. These results are discussed below.

### Similarity in survival deficit and liver damage, inflammatory markers, and bacterial accumulation

The survival rate of SR-BI and BII transgenic mice after CLP was worse compared to that of WT control mice. This finding is consistent with our earlier *in vitro* study suggesting that SR-B receptors may worsen sepsis by facilitating bacterial cytosolic accumulation, evasion of lysosomal complex, and proliferation in host cells^12^. At the same time, the survival benefit of SR-B receptors in sepsis has been also reported using SR-BI/SR-BII KO mice that showed lower survival rate after CLP than WT control. Indeed, the result was remarkably impacted by adrenal insufficiency due to impaired cholesterol ester uptake in the adrenal glands^13^. Corticosterone pretreatment (glucocorticoid only) could not reverse the survival outcome following CLP^13^. However, we reported SR-BI/BII KO had higher survival rate than WT in CLP-induced septic mice pretreated with both dexamethasone and fludrocortisone acetate^14^. As such, the survival of SR-BI/SR-BII KO after CLP varied according to the details of adrenal replacement (none: worse; glucocorticosteroids only: worse; both gluco- and mineralocorticosteroids: improved)^13, 14, 22^. Hence, we aimed to elucidate the survival impact of SR-B receptors in CLP sepsis using SR-BI and SR-BII-overexpressing mice to minimize confounding adrenal insufficiency. Our result suggests SR-BI and BII overexpression may be associated with adverse survival outcomes in sepsis.

SR-BI and BII transgenic mice had more severe histological liver damage with higher levels of liver injury markers at 24 h after CLP. Moreover, systemic inflammatory cytokines and vascular endothelial dysfunction markers are increased in SR-BI and BII transgenic mice vs control septic mice. Notably, the expression of inflammatory genes was higher in the liver of those transgenic mice, whereas the expression in kidney was similar across the experimental groups. The previous *in vitro* study exhibited the more prominent dose-dependent stimulation of cytokine production in SR-B-overexpressing HEK293 cells than mock-transfected cells across increasing doses of ligands, including *E. coli*, LPS, and GroEL^11^. Additionally, SR-BI/BII KO significantly attenuated systemic proinflammatory responses in the mice supplemented with both glucocorticoid and mineralocorticoid prior to CLP surgery. Such evidence supports profound inflammatory response and vascular endothelial impairment in septic mice overexpressing SR-BI and BII. Why was the exacerbation of the histological damage and inflammation in SR-B transgenic mice more pronounced in the liver compared to the kidney? This is probably because those mice more highly expressed and produced human SR-BI and BII in the liver and more intensive bacterial accumulation was observed in the liver based on our earlier studies^17, 18, 29^. Our results suggest that a liver hyperinflammatory state may be involved in severe liver injury and poor outcome of SR-BI and BII transgenic mice.

As we recently reported, we detected an abundance of bacteria in the liver of WT mice after CLP^29^. Unexpectedly, the bacterial count was dramatically reduced in that of SR-BI and BII transgenic mice. The immunofluorescence analyses using GFP-labeled *E. coli* demonstrated a significant increase in the frequency of bacteria localized within hepatic macrophages in SR-BI and SR-BII transgenic mice, accompanied by a reduction in the total liver bacterial number.

### Underlying cellular responses differ in SR-B1 and SR-BII transgenic mice

Despite similar suppression of liver bacteria counts in SR-BI and BII transgenic mice, the mechanisms seem to be quite different. SR-BII transgenic mice had a higher phagocytic efficiency of hepatic macrophages compared to those of SR-BI transgenic mice, consistent with our previous report showing a greater number of internalized bacteria in HeLa cells overexpressing SR-BII versus SR-BI^12^. That same study also demonstrated that bacteria internalized via SR-BI or SR-BII could survive and proliferate within host cells^12^. The discrepancy between these findings and our present results may be attributed to differences in the cell types involved in bacterial uptake.

Based on our previous *in vitro* study, BMDMs overexpressing SR-BII exhibited higher expression and production of inflammatory cytokines in response to increased phagocytosis of LPS compared to WT macrophages, whereas SR-BI-overexpressing BMDMs showed cytokine levels comparable to WT^17^. Abundant bacterial internalization may have exclusively stimulated cytokine production in SR-BII-overexpressing macrophages.

SR-BII transgenic mice also demonstrated the greatest increase in the number of F4/80-positive cells in the liver following bacterial infusion. Hepatic macrophages may have recruited circulating monocytes once activated by internalizing numerous bacteria. Hepatic macrophages were reduced in WT mice probably due to bacteria-induced Kupffer cell PANoptosis^30^.

Conversely, SR-BI transgenic mice had greater neutrophil infiltration in liver during CLP sepsis vs SR-BII mice. The pronounced neutrophil influx may be caused by partial adrenal insufficiency occurring in SR-BI (see below). Reportedly, adrenalectomy in rats facilitated the maturation of bone marrow neutrophils. Furthermore, deficiency of adrenal cortical hormones reduced L-selectin expression on polymorphonuclear (PMN) cells in the bone marrow while increasing its expression on PMN cells in systemic circulation, thereby promoting their migration into peripheral blood and subsequently to the extracellular matrix^31^.

The reduction of bacterial load was also observed in the kidney of SR-B transgenic mice, while the extent of sepsis-induced kidney damage and elevation of BUN and inflammatory cytokine expression did not differ across the experimental group. Bacteria invading from the peritoneal cavity anatomically first get trapped in the liver. Then the remaining few bacteria could reach the kidneys via the systemic circulation. The bacterial count in the kidney might have merely reflected the phagocytic activity in the liver. The bacterial accumulation in the kidney was markedly smaller than that in the liver based on our recent report^29^. Bacteria localized within F4/80-positive cells were not detected in the kidneys of any experimental group by immunofluorescence analysis. Thus, bacterial phagocytosis by macrophages may have had a limited influence on cytokine production and kidney damage during CLP-induced sepsis compared to other mechanisms, such as microcirculatory dysfunction^32, 33^.

### Difference in cholesterol metabolism and adrenal function

SR-Bs expressed on the liver and steroidogenic organs are involved in selective uptake of HDL-C^5^. SR-BI has a higher HDL-C uptake than SR-BII because SR-BI is more highly expressed on the cell surface^17^. In our study, the plasma level of HDL-C in SR-BI transgenic mice was significantly lower than that of WT at baseline and in sepsis. The result may corroborate that hepatic overexpression of SR-BI facilitated cholesterol uptake and biliary secretion, reducing cholesterol level in plasma, as in the earlier report^28^. In a dyslipidemia patient cohort, SR-BI protein levels were inversely associated with HDL-C levels^34^. Hence, the activity of SR-BI in liver may be crucial to maintain plasma HDL-C level. The Oil Red O staining demonstrated near absence of lipid droplets in the adrenal cortex of SR-BI transgenic mice at baseline. The transgenes that were driven by pLiv-11 may have been minimally expressed in adrenal glands according to the prior study^35^. Therefore, the diminished adrenal cholesterol ester storage in SR-BI transgenic mice likely reflects decreased uptake of HDL due to its low plasma level. This is similar to what is observed in SR-BI/II KO mice^36^ where diminished adrenal cholesterol ester storage is due to absent SR-BI/II which prevents cholesterol uptake despite high plasma HDL levels. Although CLP-induced septic insult increased systemic corticosterone levels in all the experimental groups, the hormone level in SR-BI transgenic mice was significantly lower than that of WT, suggesting the production of corticosterone was reduced because of the cholesterol ester shortage. In contrast, SR-BII transgenic mice exhibited only a mild reduction in corticosterone levels, probably because some amount of cholesterol ester remained in the adrenal cortex before CLP. Furthermore, replacement with both gluco- and mineralo-corticoids remarkably improved the survival of SR-BI transgenic mice to a level comparable to that of WT, whereas it had no effect on SR-BII mice. This finding demonstrates the partial adrenal insufficiency mainly contributed to the increased mortality in SR-BI mice. Not only genetic variants but also body mass index and dietary fat reportedly affect the SR-BI expression^34, 37^. Hyperglycemia and estrogen are reported to facilitate splicing shift from SR-BI to SR-BII^38, 39^. Thus, the SR-BI and BII expressions could be heterogeneous among human patients. This may explain the reason why the result of systemic steroid supplementation in human sepsis has led to conflicting results and much discussion^40^. Perhaps some of the differential responses to steroid supplementation are driven by genetic differences in the population, including SR-BI and SR-BII variants. The recent metanalysis of prospective observational studies pointed out low systemic levels of cholesterol including HDL-C at intensive care unit admission associated with hyperinflammation, high severity, and poor prognosis of critically ill patients with sepsis^41^. Further research is needed to elucidate the interplay between cholesterol metabolism, endocrine system, and immune system via SR-Bs.

## LIMITATIONS

We acknowledge several limitations of our study. Firstly, the SR-BI and SR-BII transgenic mice had human SR-B transgenes expressed not only in the liver but also in the kidney, lung, and spleen even though designed to express those receptors specifically in liver^17, 18^. The transgenes were also highly expressed in BMDMs, which can be widely distributed throughout tissues^17^. Therefore, our study could not demonstrate organ-specific human SR-B functions. Secondly, we used only male mice because the estrous cycle may affect hepatic SR-B levels; previous several reports found that estrogen regulated the expression level of SR-BI and BII in the liver^42, 43^. However, this limits the generalizability of this study. Finally, the causal relationship between bacterial phagocytosis of macrophages, adrenal insufficiency, dysregulated inflammatory response in the liver, and mortality was not completely clear in this study. Additional study is warranted to clarify the interrelationship.

In conclusion, our findings suggest human SR-BI and BII overexpression contributes to higher mortality in CLP sepsis by excessive organ damage and inflammation in the liver despite increased hepatic bacterial clearance. SR-BI transgenic mice exhibited marked neutrophil infiltration in the liver and adrenal insufficiency whereas SR-BII had prominent bacterial phagocytosis in hepatic macrophages without significant adrenal insufficiency. Thus, the increased death from sepsis may be attributable to different mechanisms in the two splice variants (Table 1). A comprehensive understanding of the pleiotropic functions of human SR-B may aid in the development of targeted therapies with reduced off-target effects. Further investigation is necessary to develop therapeutic strategies that could selectively modulate specific functions of these receptors.

**Table 1.** Possible major contributors to worsened survival in cecal ligation and puncture-induced sepsis.

| Genotype | Liver |  |  | Adrenal insufficiency |
| --- | --- | --- | --- | --- |
|  | Macrophages | Neutrophils | Inflammatory cytokines |  |
| SR-BI |  | ✓ | ✓ | partial |
| SR-BII | ✓ |  | ✓ |  |

SR-BI, class B scavenger BI; SR-BII, class B scavenger receptor BII.

## Supplementary Materials

Figure S1, Figure S2, and Figure S3.

## Author contributions

Conceptualization, NH, AVB, and PSTY.; Methodology, NH, TGV, INB, AVB, and PSTY.; Formal Analysis, NH, TGV, INB, and XH; Investigation, NH, TGV, INB, and XH.; Data Curation, NH, TGV, INB.; Writing – Original Draft Preparation, NH.; Writing – Review & Editing, TGV, INB, AVB, PSTY, APP, TLE, and RAS.; Supervision, AVB, PSTY, APP, TLE, and RAS.; Funding Acquisition, NH, TLE, and RAS.

## Institutional Review Board Statement

All the animal experiments in this study were approved by the NIDDK Animal Care and Use Committee (K100-KDB-24).

## Data Availability Statement

The datasets generated during the current study are available in the Mendeley Data repository, [https://data.mendeley.com/preview/dn35n7phbt?a=b55177cc-618a-489b-aa17-89339ca9be19].

## Acknowledgements

We would like to thank Dr. Jeffrey Reece (Advanced Light Microscopy & Image Analysis Core, National Institute of Diabetes, Digestive and Kidney Diseases, National Institutes of Health) for generously allowing us to use the confocal microscope and providing technical support.

We deeply appreciate Drs. Yoshitaka Naito (First Department of Medicine, Hamamatsu University School of Medicine, Hamamatsu, Japan) and Daiki Goto (Renal Diagnostics and Therapeutics Unit, National Institute of Diabetes, Digestive and Kidney Diseases, National Institutes of Health) for conducting randomization and blinding for this study.

## Disclaimer

This research was supported by the Intramural Research Program of the National Institute of Diabetes and Digestive and Kidney Diseases (NIDDK) within the National Institutes of Health (NIH). The contributions of the NIH author(s) were made as part of their official duties as NIH federal employees, are in compliance with agency policy requirements, and are considered Works of the United States Government. However, the findings and conclusions presented in this paper are those of the author(s) and do not necessarily reflect the views of the NIH or the U.S. Department of Health and Human Services.

---

*Supplemental Figure 1: Renal macrophages and bacteria at 8 h after bacterial injection.*

The green fluorescence protein (GFP)-labeled E. coli and macrophages were assessed with immunofluorescence analyses in kidneys harvested at 8 h after intraperitoneal bacterial infusion. The representative images for kidney specimens are exhibited (WT, n = 3; SR-BI, n = 3; SR-BII, n = 3; n = 9 total).

*Supplemental Figure 2: Green fluorescence protein-labeled E. coli in the liver and kidney at 8 h after intraperitoneal infusion.*

The immunohistochemical examination of green fluorescence protein (GFP)-labeled E. coli was performed in the liver and kidney collected from Wild-type (WT) and human class B scavenger receptor BI (SR-BI) and BII (SR-BII) transgenic mice at 8 h after injection with the bacteria. Representative images of (A) liver and (B) kidney from the experimental groups are shown (WT, n = 3; SR-BI, n = 3; SR-BII, n = 3; n = 9 total). Positive area (arrow heads) represents GFP-labeled E. coli. The bacteria were frequently detected in liver sinusoids.

## References

(1) Cajander, S.; Kox, M.; Scicluna, B. P.; Weigand, M. A.; Mora, R. A.; Flohe, S. B.; Martin-Loeches, I.; Lachmann, G.; Girardis, M.; Garcia-Salido, A.;, et al. Profiling the dysregulated immune response in sepsis: overcoming challenges to achieve the goal of precision medicine. Lancet Respir Med 2024, 12 (4), 305–322. DOI: 10.1016/S2213-2600(23)00330-2.

(2) Yuki, K.; Koutsogiannaki, S. Pattern recognition receptors as therapeutic targets for bacterial, viral and fungal sepsis. Int Immunopharmacol 2021, 98, 107909. DOI: 10.1016/j.intimp.2021.107909.

(3) Opal, S. M.; Laterre, P. F.; Francois, B.; LaRosa, S. P.; Angus, D. C.; Mira, J. P.; Wittebole, X.; Dugernier, T.; Perrotin, D.; Tidswell, M.;, et al. Effect of eritoran, an antagonist of MD2-TLR4, on mortality in patients with severe sepsis: the ACCESS randomized trial. JAMA 2013, 309 (11), 1154–1162. DOI: 10.1001/jama.2013.2194.

(4) Rice, T. W.; Wheeler, A. P.; Bernard, G. R.; Vincent, J. L.; Angus, D. C.; Aikawa, N.; Demeyer, I.; Sainati, S.; Amlot, N.; Cao, C.;, et al. A randomized, double-blind, placebo-controlled trial of TAK-242 for the treatment of severe sepsis. Crit Care Med 2010, 38 (8), 1685–1694. DOI: 10.1097/CCM.0b013e3181e7c5c9.

(5) Webb, N. R.; Connell, P. M.; Graf, G. A.; Smart, E. J.; de Villiers, W. J.; de Beer, F. C.; van der Westhuyzen, D. R. SR-BII, an isoform of the scavenger receptor BI containing an alternate cytoplasmic tail, mediates lipid transfer between high density lipoprotein and cells. J Biol Chem 1998, 273 (24), 15241–15248. DOI: 10.1074/jbc.273.24.15241.

(6) Miquel, J. F.; Moreno, M.; Amigo, L.; Molina, H.; Mardones, P.; Wistuba, II; Rigotti, A. Expression and regulation of scavenger receptor class B type I (SR-BI) in gall bladder epithelium. Gut 2003, 52 (7), 1017–1024. DOI: 10.1136/gut.52.7.1017.

(7) Zhang, Y.; Da Silva, J. R.; Reilly, M.; Billheimer, J. T.; Rothblat, G. H.; Rader, D. J. Hepatic expression of scavenger receptor class B type I (SR-BI) is a positive regulator of macrophage reverse cholesterol transport in vivo. J Clin Invest 2005, 115 (10), 2870–2874. DOI: 10.1172/JCI25327.

(8) Chinetti, G.; Gbaguidi, F. G.; Griglio, S.; Mallat, Z.; Antonucci, M.; Poulain, P.; Chapman, J.; Fruchart, J. C.; Tedgui, A.; Najib-Fruchart, J.;, et al. CLA-1/SR-BI is expressed in atherosclerotic lesion macrophages and regulated by activators of peroxisome proliferator-activated receptors. Circulation 2000, 101 (20), 2411–2417. DOI: 10.1161/01.cir.101.20.2411.

(9) Brundert, M.; Ewert, A.; Heeren, J.; Behrendt, B.; Ramakrishnan, R.; Greten, H.; Merkel, M.; Rinninger, F. Scavenger receptor class B type I mediates the selective uptake of high-density lipoprotein-associated cholesteryl ester by the liver in mice. Arterioscler Thromb Vasc Biol 2005, 25 (1), 143–148. DOI: 10.1161/01.ATV.0000149381.16166.c6.

(10) Ji, Y.; Jian, B.; Wang, N.; Sun, Y.; Moya, M. L.; Phillips, M. C.; Rothblat, G. H.; Swaney, J. B.; Tall, A. R. Scavenger receptor BI promotes high density lipoprotein-mediated cellular cholesterol efflux. J Biol Chem 1997, 272 (34), 20982–20985. DOI: 10.1074/jbc.272.34.20982.

(11) Baranova, I. N.; Vishnyakova, T. G.; Bocharov, A. V.; Leelahavanichkul, A.; Kurlander, R.; Chen, Z.; Souza, A. C.; Yuen, P. S.; Star, R. A.; Csako, G.;, et al. Class B scavenger receptor types I and II and CD36 mediate bacterial recognition and proinflammatory signaling induced by Escherichia coli, lipopolysaccharide, and cytosolic chaperonin 60. J Immunol 2012, 188 (3), 1371–1380. DOI: 10.4049/jimmunol.1100350.

(12) Vishnyakova, T. G.; Kurlander, R.; Bocharov, A. V.; Baranova, I. N.; Chen, Z.; Abu-Asab, M. S.; Tsokos, M.; Malide, D.; Basso, F.; Remaley, A.;, et al. CLA-1 and its splicing variant CLA-2 mediate bacterial adhesion and cytosolic bacterial invasion in mammalian cells. Proc Natl Acad Sci U S A 2006, 103 (45), 16888–16893. DOI: 10.1073/pnas.0602126103.

(13) Guo, L.; Song, Z.; Li, M.; Wu, Q.; Wang, D.; Feng, H.; Bernard, P.; Daugherty, A.; Huang, B.; Li, X. A. Scavenger Receptor BI Protects against Septic Death through Its Role in Modulating Inflammatory Response. J Biol Chem 2009, 284 (30), 19826–19834. DOI: 10.1074/jbc.M109.020933.

(14) Leelahavanichkul, A.; Bocharov, A. V.; Kurlander, R.; Baranova, I. N.; Vishnyakova, T. G.; Souza, A. C.; Hu, X.; Doi, K.; Vaisman, B.; Amar, M.;, et al. Class B scavenger receptor types I and II and CD36 targeting improves sepsis survival and acute outcomes in mice. J Immunol 2012, 188 (6), 2749–2758. DOI: 10.4049/jimmunol.1003445.

(15) Catanese, M. T.; Ansuini, H.; Graziani, R.; Huby, T.; Moreau, M.; Ball, J. K.; Paonessa, G.; Rice, C. M.; Cortese, R.; Vitelli, A.;, et al. Role of scavenger receptor class B type I in hepatitis C virus entry: kinetics and molecular determinants. J Virol 2010, 84 (1), 34–43. DOI: 10.1128/JVI.02199-08.

(16) Gowda, N. M.; Wu, X.; Kumar, S.; Febbraio, M.; Gowda, D. C. CD36 contributes to malaria parasite-induced pro-inflammatory cytokine production and NK and T cell activation by dendritic cells. PLoS One 2013, 8 (10), e77604. DOI: 10.1371/journal.pone.0077604.

(17) Baranova, I. N.; Souza, A. C.; Bocharov, A. V.; Vishnyakova, T. G.; Hu, X.; Vaisman, B. L.; Amar, M. J.; Chen, Z.; Kost, Y.; Remaley, A. T.;, et al. Human SR-BI and SR-BII Potentiate Lipopolysaccharide-Induced Inflammation and Acute Liver and Kidney Injury in Mice. J Immunol 2016, 196 (7), 3135–3147. DOI: 10.4049/jimmunol.1501709.

(18) Baranova, I. N.; Bocharov, A. V.; Vishnyakova, T. G.; Chen, Z.; Birukova, A. A.; Ke, Y.; Hu, X.; Yuen, P. S. T.; Star, R. A.; Birukov, K. G.;, et al. Class B Scavenger Receptors BI and BII Protect against LPS-Induced Acute Lung Injury in Mice by Mediating LPS. Infect Immun 2021, 89 (10), e0030121. DOI: 10.1128/IAI.00301-21.

(19) Simonet, W. S.; Bucay, N.; Lauer, S. J.; Taylor, J. M. A far-downstream hepatocyte-specific control region directs expression of the linked human apolipoprotein E and C-I genes in transgenic mice. J Biol Chem 1993, 268 (11), 8221–8229.

(20) Tsuji, N.; Tsuji, T.; Yamashita, T.; Hayase, N.; Hu, X.; Yuen, P. S.; Star, R. A. BAM15 treats mouse sepsis and kidney injury, linking mortality, mitochondrial DNA, tubule damage, and neutrophils. J Clin Invest 2023, 133 (7). DOI: 10.1172/JCI152401.

(21) Shrum, B.; Anantha, R. V.; Xu, S. X.; Donnelly, M.; Haeryfar, S. M.; McCormick, J. K.; Mele, T. A robust scoring system to evaluate sepsis severity in an animal model. BMC Res Notes 2014, 7, 233. DOI: 10.1186/1756-0500-7-233.

(22) Ai, J.; Guo, L.; Zheng, Z.; Wang, S. X.; Huang, B.; Li, X. A. Corticosteroid Therapy Benefits Septic Mice With Adrenal Insufficiency But Harms Septic Mice Without Adrenal Insufficiency. Crit Care Med 2015, 43 (11), e490–498. DOI: 10.1097/CCM.0000000000001264.

(23) Tiwari, M. M.; Messer, K. J.; Mayeux, P. R. Inducible nitric oxide synthase and apoptosis in murine proximal tubule epithelial cells. Toxicol Sci 2006, 91 (2), 493–500. DOI: 10.1093/toxsci/kfj168.

(24) Scott, M. J.; Liu, S.; Shapiro, R. A.; Vodovotz, Y.; Billiar, T. R. Endotoxin uptake in mouse liver is blocked by endotoxin pretreatment through a suppressor of cytokine signaling-1-dependent mechanism. Hepatology 2009, 49 (5), 1695–1708. DOI: 10.1002/hep.22839.

(25) Aderhold, A.; Alexaki, V. I. Lipid metabolism in the adrenal gland. Front Endocrinol (Lausanne) 2025, 16, 1577505. DOI: 10.3389/fendo.2025.1577505.

(26) Cai, L.; Ji, A.; de Beer, F. C.; Tannock, L. R.; van der Westhuyzen, D. R. SR-BI protects against endotoxemia in mice through its roles in glucocorticoid production and hepatic clearance. J Clin Invest 2008, 118 (1), 364–375. DOI: 10.1172/JCI31539.

(27) Trigatti, B.; Rayburn, H.; Vinals, M.; Braun, A.; Miettinen, H.; Penman, M.; Hertz, M.; Schrenzel, M.; Amigo, L.; Rigotti, A.;, et al. Influence of the high density lipoprotein receptor SR-BI on reproductive and cardiovascular pathophysiology. Proc Natl Acad Sci U S A 1999, 96 (16), 9322–9327. DOI: 10.1073/pnas.96.16.9322.

(28) Yesilaltay, A.; Morales, M. G.; Amigo, L.; Zanlungo, S.; Rigotti, A.; Karackattu, S. L.; Donahee, M. H.; Kozarsky, K. F.; Krieger, M. Effects of hepatic expression of the high-density lipoprotein receptor SR-BI on lipoprotein metabolism and female fertility. Endocrinology 2006, 147 (4), 1577–1588. DOI: 10.1210/en.2005-1286.

(29) Hayase, N.; Chari, R. R.; Dos Santos, A. A. C.; Naito, Y.; Hu, X.; Yuen, P. S. T.; Star, R. A. Continuous antithrombin III infusion in a clinically relevant sepsis model. Shock 2025. DOI: 10.1097/SHK.0000000000002648.

(30) Li, T.; Adams, J.; Zhu, P.; Zhang, T.; Tu, F.; Gravitte, A.; Zhang, X.; Liu, L.; Casteel, J.; Yakubenko, V.;, et al. The role of heme in sepsis induced Kupffer cell PANoptosis and senescence. Cell Death Dis 2025, 16 (1), 284. DOI: 10.1038/s41419-025-07637-6.

(31) Cavalcanti, D. M.; Lotufo, C. M.; Borelli, P.; Tavassi, A. M.; Pereira, A. L.; Markus, R. P.; Farsky, S. H. Adrenal deficiency alters mechanisms of neutrophil mobilization. Mol Cell Endocrinol 2006, 249 (1-2), 32–39. DOI: 10.1016/j.mce.2006.01.007.

(32) Ergin, B.; Kapucu, A.; Demirci-Tansel, C.; Ince, C. The renal microcirculation in sepsis. Nephrol Dial Transplant 2015, 30 (2), 169–177. DOI: 10.1093/ndt/gfu105.

(33) Betrie, A. H.; Ma, S.; Ow, C. P. C.; Peiris, R. M.; Evans, R. G.; Ayton, S.; Lane, D. J. R.; Southon, A.; Bailey, S. R.; Bellomo, R.;, et al. Renal arterial infusion of tempol prevents medullary hypoperfusion, hypoxia, and acute kidney injury in ovine Gram-negative sepsis. Acta Physiol (Oxf) 2023, 239 (1), e14025. DOI: 10.1111/apha.14025.

(34) West, M.; Greason, E.; Kolmakova, A.; Jahangiri, A.; Asztalos, B.; Pollin, T. I.; Rodriguez, A. Scavenger receptor class B type I protein as an independent predictor of high-density lipoprotein cholesterol levels in subjects with hyperalphalipoproteinemia. J Clin Endocrinol Metab 2009, 94 (4), 1451–1457. DOI: 10.1210/jc.2008-1223 From NLM Medline.

(35) Temel, R. E.; Tang, W.; Ma, Y.; Rudel, L. L.; Willingham, M. C.; Ioannou, Y. A.; Davies, J. P.; Nilsson, L. M.; Yu, L. Hepatic Niemann-Pick C1-like 1 regulates biliary cholesterol concentration and is a target of ezetimibe. J Clin Invest 2007, 117 (7), 1968–1978. DOI: 10.1172/JCI30060.

(36) Ito, M.; Ye, X.; Wang, Q.; Guo, L.; Hao, D.; Howatt, D.; Daugherty, A.; Cai, L.; Temel, R.; Li, X. A. SR-BI (Scavenger Receptor BI), Not LDL (Low-Density Lipoprotein) Receptor, Mediates Adrenal Stress Response-Brief Report. Arterioscler Thromb Vasc Biol 2020, 40 (8), 1830–1837. DOI: 10.1161/ATVBAHA.120.314506 From NLM Medline.

(37) Spady, D. K.; Kearney, D. M.; Hobbs, H. H. Polyunsaturated fatty acids up-regulate hepatic scavenger receptor B1 (SR-BI) expression and HDL cholesteryl ester uptake in the hamster. J Lipid Res 1999, 40 (8), 1384–1394. From NLM Medline.

(38) Chiba-Falek, O.; Nichols, M.; Suchindran, S.; Guyton, J.; Ginsburg, G. S.; Barrett-Connor, E.; McCarthy, J. J. Impact of gene variants on sex-specific regulation of human Scavenger receptor class B type 1 (SR-BI) expression in liver and association with lipid levels in a population-based study. BMC Med Genet 2010, 11, 9. DOI: 10.1186/1471-2350-11-9 From NLM Medline.

(39) Mendoza, G. V. S., Susana Elfrida; González, Irma Inés; Della Vedova, Maria CeciliaIcon; Fernandez, Gustavo; Ojeda, Marta Susana. Hyperglycemia promotes overexpression of SR-BII isoform of the scavenger receptor class B type I in type 2 diabetes mellitus: A study in Juana Koslay City, San Luis, Argentina. Journal of Diabetes Mellitus 2013, 3 (4), 172–183. DOI: 10.4236/jdm.2013.34027.

(40) Patel, G. P.; Balk, R. A. Systemic steroids in severe sepsis and septic shock. Am J Respir Crit Care Med 2012, 185 (2), 133–139. DOI: 10.1164/rccm.201011-1897CI From NLM Medline.

(41) Taylor, R.; Zhang, C.; George, D.; Kotecha, S.; Abdelghaffar, M.; Forster, T.; Santos Rodrigues, P. D.; Reisinger, A. C.; White, D.; Hamilton, F.;, et al. Low circulatory levels of total cholesterol, HDL-C and LDL-C are associated with death of patients with sepsis and critical illness: systematic review, meta-analysis, and perspective of observational studies. EBioMedicine 2024, 100, 104981. DOI: 10.1016/j.ebiom.2024.104981.

(42) Stangl, H.; Graf, G. A.; Yu, L.; Cao, G.; Wyne, K. Effect of estrogen on scavenger receptor BI expression in the rat. J Endocrinol 2002, 175 (3), 663–672. DOI: 10.1677/joe.0.1750663.

(43) Zhang, X.; Moor, A. N.; Merkler, K. A.; Liu, Q.; McLean, M. P. Regulation of alternative splicing of liver scavenger receptor class B gene by estrogen and the involved regulatory splicing factors. Endocrinology 2007, 148 (11), 5295–5304. DOI: 10.1210/en.2007-0376.

