## Supplementary figures and images for "Different functions of human scavenger receptors BI and BII overexpressed in a murine abdominal sepsis model"

### Supplemental Figure 1

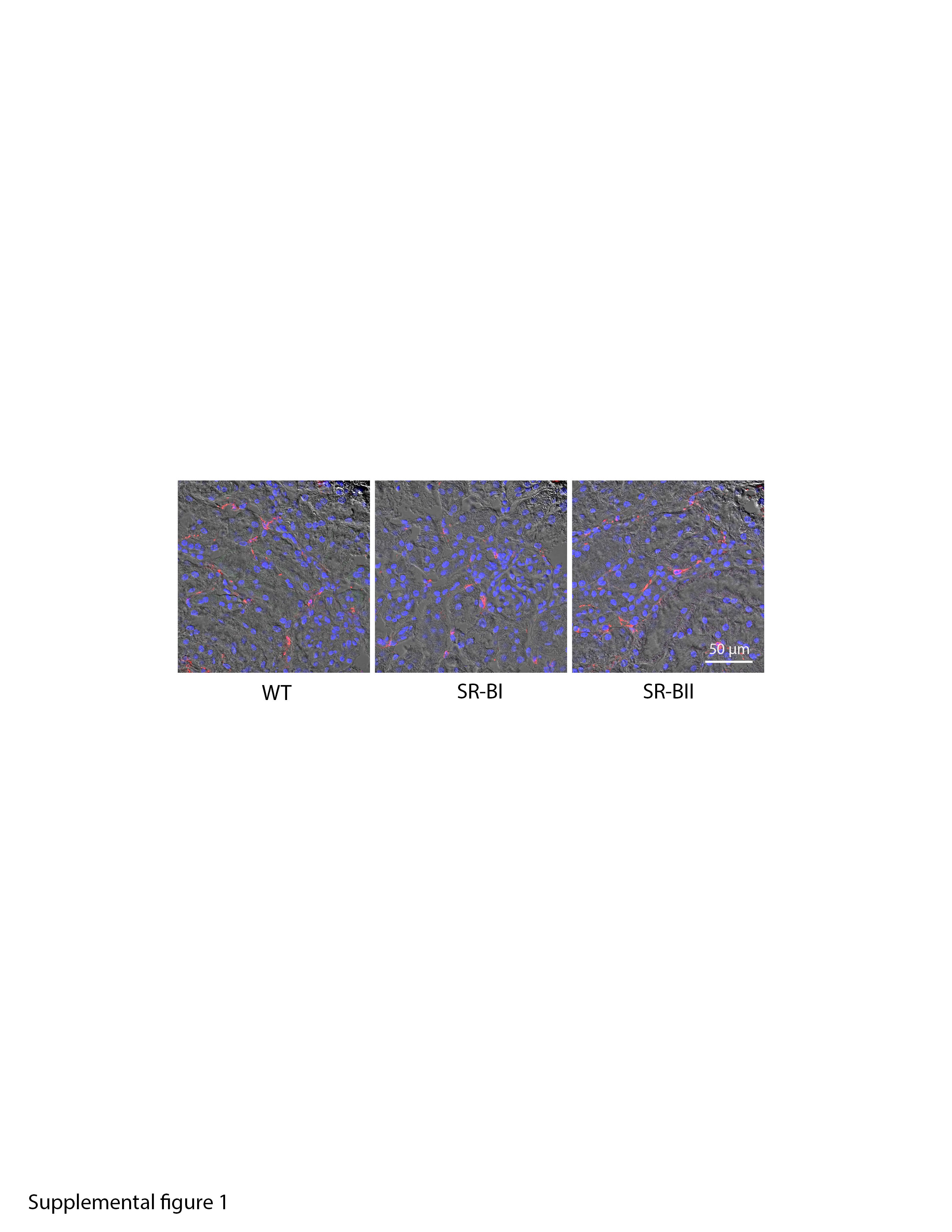

### Supplemental Figure 2

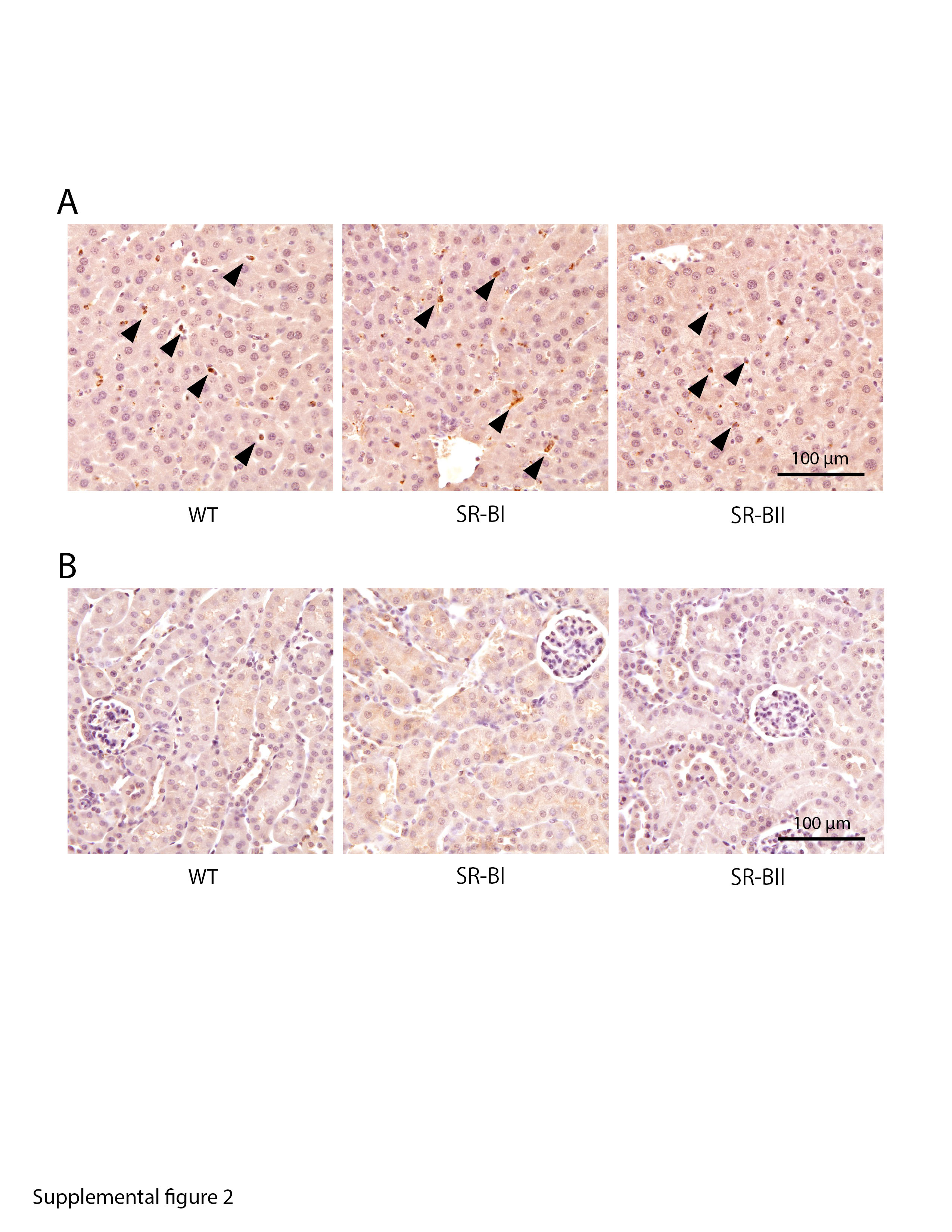
